# Flexible and accurate decoding of neural populations through stochastic comodulation

**DOI:** 10.1101/624387

**Authors:** Caroline Haimerl, Cristina Savin, Eero P. Simoncelli

## Abstract

Sensory-guided behavior requires reliable encoding of information (from stimuli to neural responses) and flexible decoding (from neural responses to behavior). In typical decision tasks, a small subset of cells within a large population encode task-relevant stimulus information and need to be *identified* by later processing stages for relevant information to be transmitted. A statistically optimal decoder (e.g., maximum likelihood) can utilize task-relevant cells for any given task configuration, but relies on complete knowledge of the relationship between the task and the stimulus-response and noise properties of the encoding population. The brain could learn an optimal decoder for a task through supervised learning (i.e., regression), but this typically requires many training trials, and thus lacks the flexibility of humans or animals, that can rapidly adjust to changes in task parameters or structure. Here, we propose a novel decoding solution based on functionally targeted stochastic modulation. Population recordings during different discrimination tasks have revealed that a substantial portion of trial-to-trial variability in cell responses can be explained by stochastic modulatory signals that are shared, and that seem to preferentially target task-informative neurons (Rabinowitz et al., 2015). The variability introduced by these modulators corrupts the encoded stimulus signal, but we propose that it also serves as a label for the informative neurons, allowing the decoder to solve the identification problem. We show in simulations of a modulated Poisson spiking model that a linear decoder with readout weights proportional to the estimated neuron-specific strength of modulation achieves performance close to an optimal decoder.

## 1 Introduction

Our survival depends on the actions we take, which are derived from our internal state and sensory input. Specifically, accurate decisions require reliable encoding and flexible task-specific decoding of sensory information. Take for instance the perceptual task of detecting a change in orientation of a grating within a small aperture, placed at a particular location in the visual field (Fig. 1). Neurons in primary visual cortex (V1) that respond selectively to edges at different spatial locations and orientations encode the visual stimulus. However, only a small fraction of those neurons would show a change in response when the stimulus changes orientation in the task (Fig. 1, red); the overwhelming majority will not respond at all or their responses would not vary as the stimulus changes. Since nearly all visual information passes through V1, a downstream areas’ sole source of information lies in the responses of those few V1 cells. Thus, solving this task relies on the ability to properly gather and combine the responses of these task-relevant neurons, while ignoring the background chatter of activity emanating from the remainder of the population. Furthermore, if the task changes (e.g. through a change in stimulus position or orientation), the informative sub-population within V1 will change, and downstream areas will need to flexibly modify their processing. It remains a mystery how such dynamic task-dependent routing of information is achieved in the brain.

**Figure 1:**
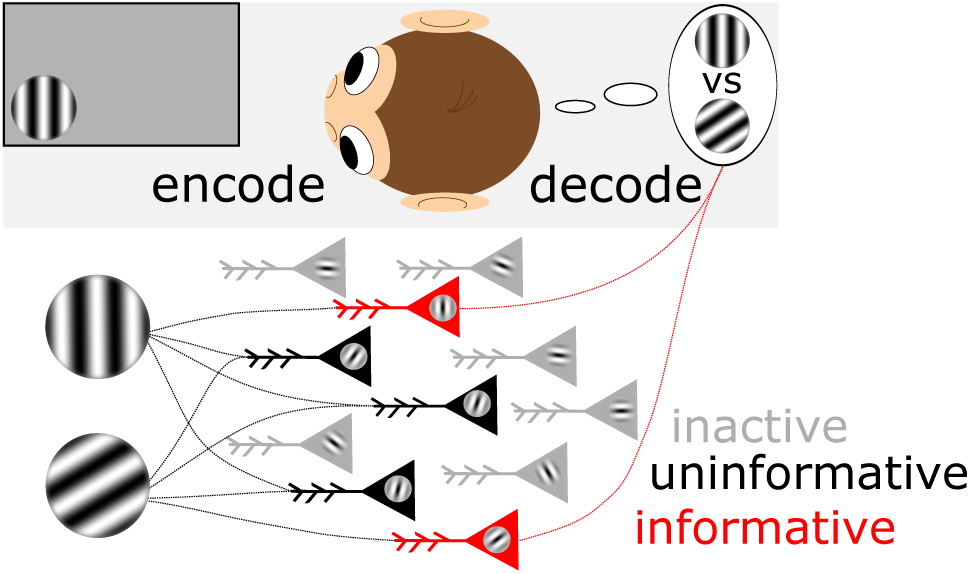
Illustration of the encoding-decoding framework. In a binary fine discrimination task with two similar fixed stimuli (here oriented gratings), a subpopulation of encoding neurons will respond strongly to the stimuli while most neurons will be respond weakly or not at all (grey). Among the subpopulation of active neurons, the subset that are activated primarily by one of the two stimuli (red) carry information relevant for the discrimination of the two stimuli, while the remaining active neurons are uninformative (black). A decoding area needs to pay attention to the informative neurons (while downplaying the others), and read out the sensory signal embedded in their noisy activity.

The readout of sensory information in neural responses is often explored using statistically optimal decoders derived from specific encoding models, which set an upper bound on performance (Shadlen et al., 1996; Seung and Sompolinsky, 1993; Salinas and Abbott, 1994; Ma et al., 2006; Jazayeri and Movshon, 2007; Caruana, 1997; Britten et al., 1996; Ganguli and Simoncelli, 2016). Commonly these decoders rely on full knowledge of the stimulus response and noise properties of neurons, so as to weigh their individual contributions optimally. These assumptions make up the *ideal observer* framework (Dayan and Abbott, 2005; Geisler and Albrecht, 1997), which takes the view of a scientist who records cellular activity during perception over many controlled trials). However, it seems inconceivable that upstream decoding circuits could store or have access to such detailed information. The alternative is to learn the decoder from experience. This however requires extensive training on the discrimination task, accompanied with feedback about success or failure on each trial. Such slow learning seems inconsistent with the observed behavioral flexibility of animals and humans, both of whom can rapidly adjust to changes in task conditions (Simoncelli, 2009).

Our understanding of sensory encoding is better developed, and instantiated in models that explain substantial portions of stimulus-driven response, as well as variability. This variability is usually treated as noise (Shadlen et al., 1996) which limits the amount of stimulus information that a neural population can encode (Averbeck et al., 2006). While noise in neuron’s activity is commonly modeled with a Poisson process, the trial-to-trial variability of a population given a constant stimulus often exceeds the expected variance under a Poisson process, suggesting that additional sources of time-varying modulation influence the response of neurons (Goris et al., 2014). This modulation was found to be low-dimensional (Rabinowitz et al., 2015) (i.e., shared across neurons), reflected by varying degrees of pairwise noise correlation (Cohen and Maunsell, 2009; Ruff et al., 2016; Ni et al., 2017; Bondy et al., 2018). Moreover, it is primarily multiplicative, stochastic and differing in strength across neurons (Rabinowitz et al., 2015; Franke et al., 2016; Lin et al., 2015). Theoretical work indicates that such correlated noise can be detrimental for population encoding, as it cannot be averaged out (Britten et al., 1996; Moreno-Bote et al., 2014). Even worse, in some experiments this noise seems to be specifically targeted to neurons that are informative for the task (Rabinowitz et al., 2015; Bondy et al., 2018), further exacerbating these detrimental effects.

While correlated noise places limitations on the brain’s ability to encode information accurately, here we show that it may serve as key ingredient in solving the mystery of flexible decoding. We put forward a theoretical framework based on an observation of the properties of an inferred low-dimensional modulator: (Rabinowitz et al., 2015) analyzed V4 population data by (Cohen and Maunsell, 2009), and found that neurons that are informative for the task were more strongly modulated (see Fig. 2), making them also more strongly correlated. Similarly, (Bondy et al., 2018) found that V1 noise correlation structure is better explained by task-informativeness than by stimulus tuning properties, suggesting that the source of this correlation is a top down (as opposed to stimulus-driven) signal. These results are counter-intuitive as they suggest that a noisy task-irrelevant modulator interferes with the responses of task-informative neurons, thereby degrading the encoded stimulus information. While the origin of this noise-introducing modulator remains unclear, we propose that its negative impact on stimulus-encoding comes with an essential compensatory decoding benefit: the modulatory fluctuations serve as a “label” for the task-relevant neurons, helping the decoder to select these neurons for readout. The proposed scheme requires that the decoder make use of the modulator itself or the modulator-induced covariability, when assigning appropriate decoding weights to each neuron. We develop such a *modulator-guided decoder*, and show through simulations that moderate levels of task-specific stochastic modulation of an encoding population can lead to a substantial overall benefit in decoding accuracy, while keeping the assumed knowledge about the encoding population at a biologically plausible level. So rather than being a bug, structured noise may be an essential feature of brain computation.

**Figure 2:**
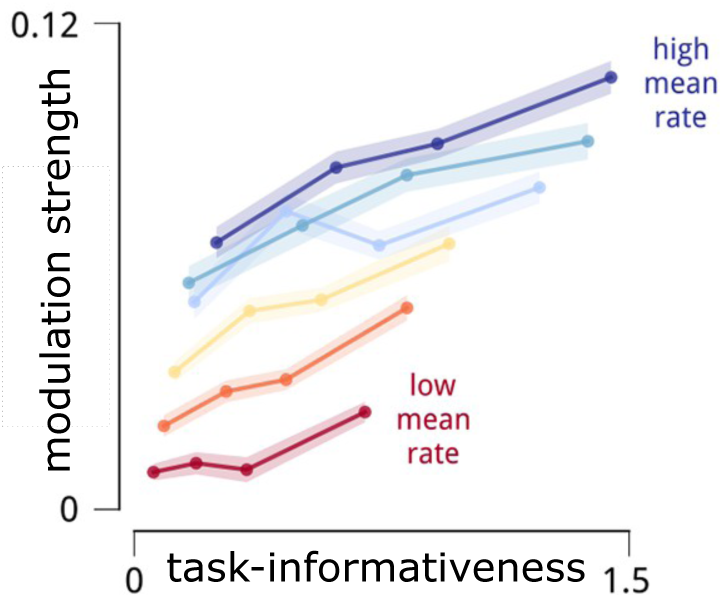
Experimental evidence for modulatory labelling (modified from Rabinowitz et al. (2015)). During a visual discrimination task, V4 neurons are modulated by a shared stochastic signal. The strength of this modulation is highly correlated with their task-informativeness, even when restricted to subsets of neurons with similar firing rates.

## 2 Encoding/decoding models

To test our hypothesis, we simulate encoding in a population of stimulus-selective, noise-modulated Poisson neurons (Rabinowitz et al., 2015) and compare statistically optimal *ideal observer* decoders, that have full knowledge of the stimulus-selectivity and modulatory structure of the encoding population, with *biologically plausible* decoders, that have limited knowledge of the encoding population.

### Encoding model: Poisson spiking population with task-targeted modulation

The variability in spike count response *k*_*t*_ over repeated presentations of a stimulus *s* reflects the stochastic nature of neural spiking, commonly modeled using a Poisson point process with stimulus-dependent firing rate *λ*(*s*). We account for supra-Poisson variability in neural responses, by introducing additional sources of stochasticity (Goris et al., 2014; Lin et al., 2015). Specifically, the stimulus-driven rate is dynamically modulated by a time-varying signal *m*_*t*_ (Rabinowitz et al., 2015), which leads to a doubly stochastic spiking process:

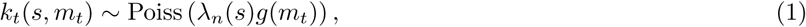

where *g*(·) is a positive-valued link function to guarantee a positive firing rate, here an exponential.

We simulate a binary discrimination task (i.e., discriminate *s* = 0 from *s* = 1) similar to the change-detection task used by Cohen and Maunsell (2009). Empirical observations in macaque area V4 show that the modulatory signal *m*_*t*_ is low-dimensional, shared across the neural population, and selectively targets neurons proportional to their task-informativeness (Rabinowitz et al., 2015). We therefore introduce neuron-specific modulation strength, *w*_*n*_, that is correlated with the *n*th neuron’s ability to discriminate the two stimuli:

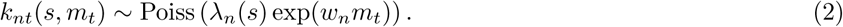

Following experimental results (Rabinowitz et al., 2015), we use i.i.d. zero-mean Gaussian noise and variance 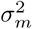 for *m*_*t*_. In simulations we simplify further by using binary modulation weights (i.e., only task-informative neurons are modulated).

### Statistically optimal “ideal observer” decoders

Given the modulated Poisson encoding model, an ideal observer with complete knowledge of both stimulus response properties {*λ*_*n*_(*s*)} and modulation properties {*w*_*n*_, *m*_*t*_} provides an upper-bound on task-decision accuracy. It does so by comparing the probability of the two stimuli under the full model, or equivalently, examining the sign of their log odds. For our modulated Poisson encoding model (see Eq. 2), this reduces optimal discrimination to comparing a weighted linear combination of the observed neural spike counts against a time-varying threshold that is a function of the modulator (see details in Appendix). We refer to it as the *modulator-conditioned maximum likelihood* (MC-ML) decoder^1^:

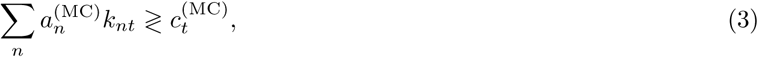

with optimal weights:

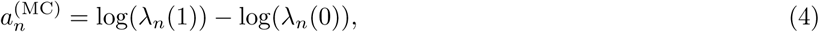

and *time-varying* threshold:

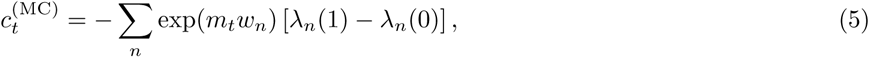

where *λ*_*n*_(*s*) denotes the mean response of the *n*-th neuron to stimulus *s* for *m*_*t*_ = 0.

The MC-ML decoder provides an upper bound on achievable performance, and relies on perfect knowledge of the modulator *m*_*t*_, the stimulus selectivity of the cells, *λ*_*n*_(*s*), and the coupling weights *w*_*n*_. We can relax these requirements, by assuming that the modulator is unknown, and only the modulator-marginalized stimulus selectivity of the cells is available (i.e., the stimulus response averaged over the modulator - see Appendix). We refer to this solution as the *modulator-marginalized maximum likelihood* (MM-ML) decoder.

Due to the particularities of the Poisson noise model, this second decoder also computes a weighted sum over responses:

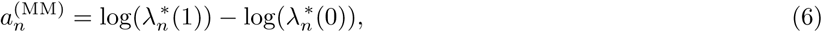

and compares this to a *fixed* threshold:

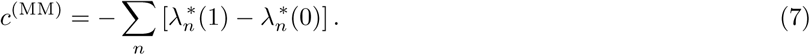

Here,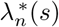 is the mean response of the *n*th neuron averaged (marginalized) over possible modulator values:

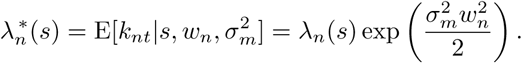

Since the exponential modulation factor cancels out when subtracting the likelihoods of the two stimuli, this yields the same decoding weights as used by the MC-ML decoder (i.e.,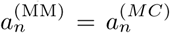, see Appendix A1). Hence, for a binary discrimination task, estimating decoding weights from the stimulus responses, without measuring modulation, still achieves an unbiased estimate. However, it does lead to systematic biases in the decoder threshold.

### Biologically plausible decoders

The MC-ML and MM-ML decoders do not seem plausible as a description of decoding in the brain, but they do provide a useful yardstick against which to compare the performance of more realistic decoders. They also indicate the use of a linear-threshold functional form for the solution. We now seek decoders of this form, that satisfy three criteria: (1) they are biologically plausible, in that they do not rely on detailed *knowledge* about the encoding population (neither the stimulus responses, nor the modulation weights), (2) they are behaviorally plausible, in that they have the ability to efficiently *adapt* to changes in task structure, so as to reflect the flexibility seen in monkey behavior (Cohen and Maunsell, 2009; Simoncelli, 2009), and (3) they achieve *accuracy* approaching that of the optimal decoders.

We start with the simplest instantiation of an approximate decoder, motivated by early work on neural binary discrimination/detection (Shadlen et al., 1996). The idea is to average the response of two sub-populations (“preferred” and “anti-preferred”) and then compare these averages. Hence, the problem of learning decoding weights is reduced to choosing which population each neuron is assigned to; this is mathematically equivalent to determining the signs of a unit weight vector (with all weights having equal magnitude). For this reason, we refer to this model as the *sign-only* (SO) decoder. The signs can be estimated via a simple learning rule, by comparing the mean response to one stimulus versus the other. Moreover, this solution is agnostic to the details of the encoding model.

The SO decoder assumes minimal knowledge of the encoding population, satisfying our first criterion. In order for this decoders to satisfy our second criterion – decoding flexibility – learning the signs should not take many trials. Indeed we see that classification into the two signed groups reaches high (90%) accuracy with only a few 10s of trials, assuming low to moderate modulator strength (see Fig. 5). If all neurons in a population were informative, learning the signs would provide an accurate readout of task information and the SO decoder would successfully fulfill also the last criterion (decoding accuracy). However, neural populations are diverse, and would generally be expected to include many noninformative neurons. For example, (Cohen and Newsome, 2009; Britten et al., 1996) find that about ∼10 – 20% of the measured, active neurons are also informative. The exact percentage depends on the neural population and behavioral task (see Appendix A4). An illustration is given in Fig. 4 which shows responses of a simulated population of encoding neurons to two task-specific stimuli. Only a small fraction of neurons are responsive, while the large majority of neurons respond weakly (“inactive”).

**Figure 3:**
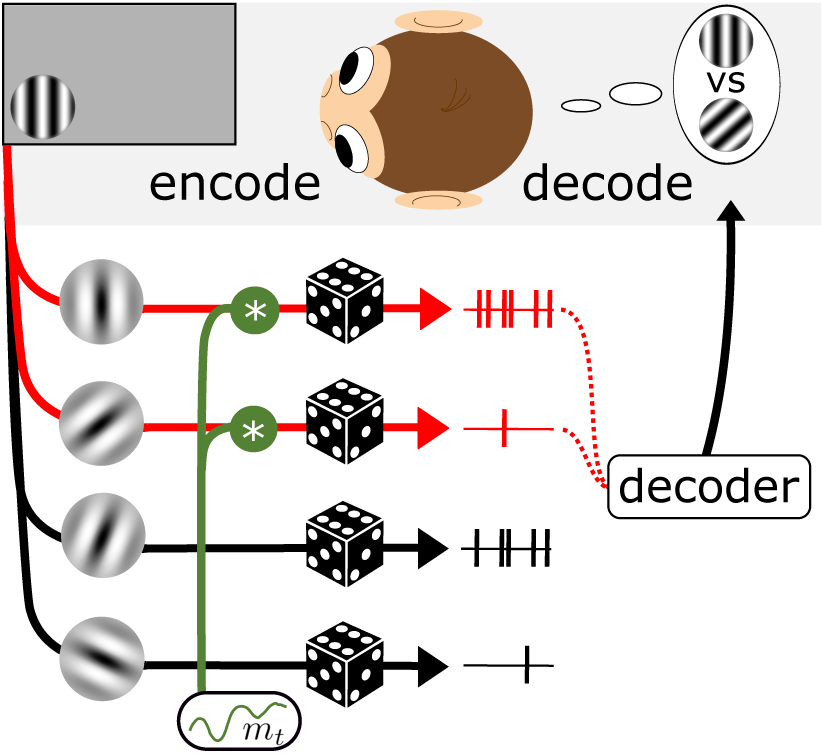
Encoding-decoding model with stochastic modulation. An encoding population consists of Poisson spiking neurons. Each neuron’s firing rate is set by its stimulus response properties. Additionally, informative neurons receive shared, stochastic modulation (green). A decoder then uses the modulatory label to identify these task-informative neuron, and uses them to arrive at the decision.

**Figure 4:**
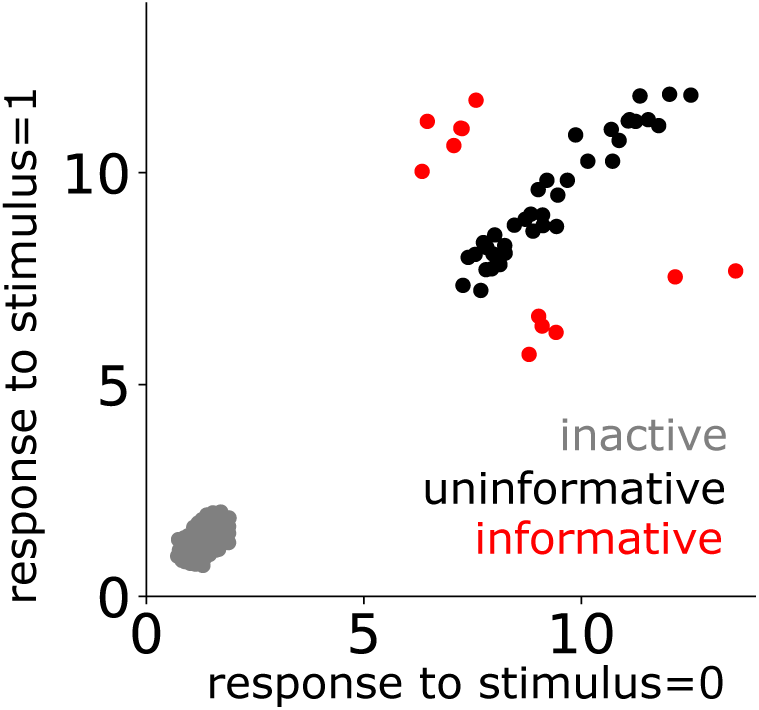
Example simulated population, as used in our simulations. Neurons fall into three categories, based on their mean response to each of the two stimuli in the discrimination task. Neurons that respond differently to the two stimuli are informative (red). Neurons with substantial, but nearly equal, responses to each stimulus are uninformative (black). The remaining neurons are inactive, showing weak responses to both stimuli (gray).

**Figure 5:**
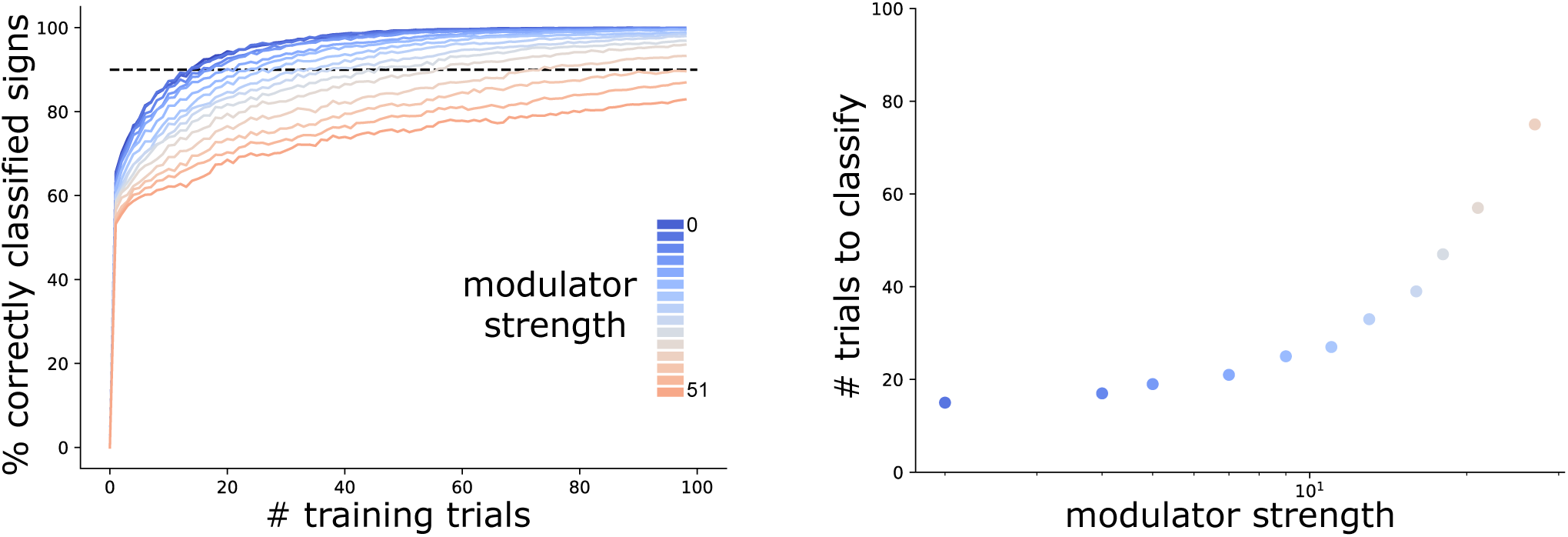
Accuracy of sign estimation, for simulated data. Left; Mean % correctly attributed signs as a function of number of training trials with varying modulator strengths (modulator makes up between 0 –51% of the spike count variance of the informative neurons). Decoding signs are learned within a few tens of trials. Right; Mean number of trials required for 90% classification accuracy, as a function of modulator strength.

If the noise from these inactive neurons is not excluded by the decoder it can corrupt the signal (Shadlen et al., 1996). We assessed decoding performance (% accurate) as a function of the number of inactive neurons (Fig. 6). The SO decoder includes inactive neurons and assigns them to one decoding group or the other based on noise alone, so that the task-irrelevant responses eventually dominate the relevant stimulus signal (see Fig. 6). In order to discount the inactive neurons, they should be assigned decoding weights with smaller amplitudes. The limited knowledge constraint, would however mean that these weights cannot be assumed to be known, but must be learned/adapted based on information readily available to the decoder. Since informative neurons necessarily have to show activity during a task, one simple, heuristic rule is to set decoding weights proportional to the mean spike count of their corresponding neurons:

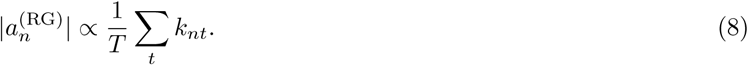

**Figure 6:**
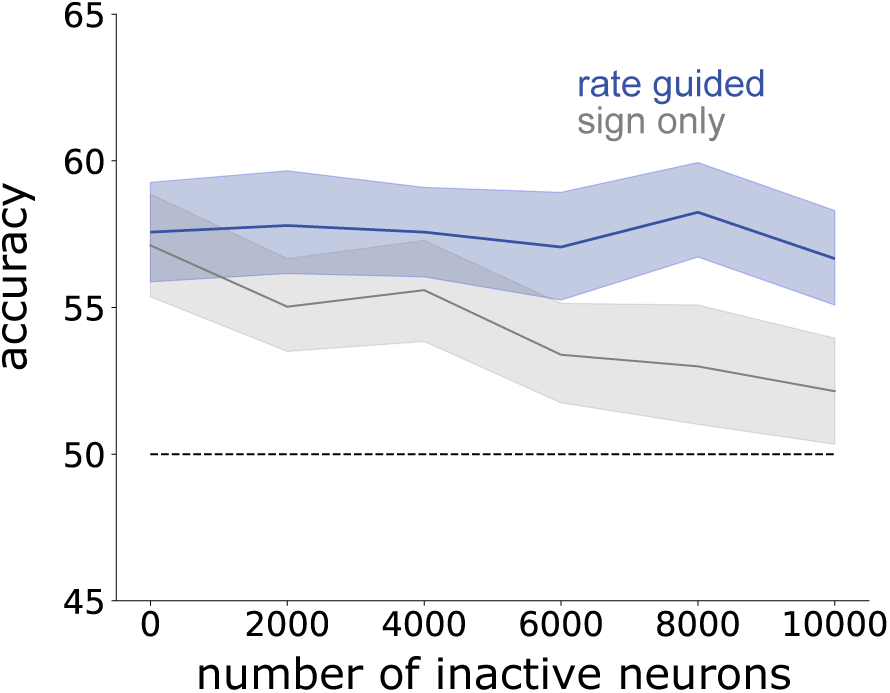
Mean performance of RG and SO decoders as the number of inactive neurons is increased. The RG decoder downweights inactive neurons, thus allowing it to maintain performance.

For this decoder, the sign of the weights must again be learned (as for the SO decoder). The threshold is set optimally but time-invariant. This *rate-guided* (RG) decoder improves decoding accuracy over the SO decoder by excluding neurons that do not respond to the stimuli (grey in Fig. 4). We see that while the SO decoder’s performance drops to chance level with increasing numbers of inactive neurons, the RG decoder is unaffected. However, the RG decoder still does not achieve a behavioral level of performance (*∼* 90% accuracy). This is because it can not exclude neurons that are active, but respond similarly to both stimuli (“uninformative neurons”, Fig. 4).

To overcome this limitation, we developed a *modulator-guided* (MG) decoder that exploits the task-specific modulatory structure of the encoding model to set decoding weights. We assume the modulator *m*_*t*_ is known (e.g., as a broadcast top-down signal – see Discussion), and estimate the neuron-specific modulation strength by computing the inner product between each neuron’s response and the modulator, thereby obtaining also an estimate of optimal decoding weights:

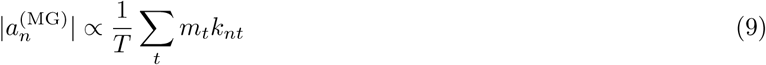

As for the simpler decoders, the signs of the decoding weights can be estimated via the learning rule of the SO decoder. The modulator-dependent threshold has the same form as the time-varying threshold defined by the MC-ML decoder, but the true modulation weights *w*_*n*_ must be replaced by empirical estimates:

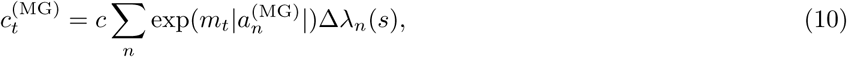

and the difference in firing rates [*λ*_*n*_(1) – *λ*_*n*_(0)] is replaced by an estimate Δ*λ*_*n*_(*s*) that is a function of the estimated decoding weights, the learned signs and one free parameter per informative subpopulation (two parameters in total). This parameter approximates the average change in activity as the stimulus changes and can easily be learned (see details in Appendix A5).

## 3 Decoder accuracy

We tested the decoders in Table 1, in a binary discrimination task that evokes differential responses in a small subset of cells in the encoding population. Decoding performance (e.g., estimation of *s*, or discrimination between two values of *s*) depends on the strength of the modulation. Specifically, the modulator variance controls overall modulation strength in the population (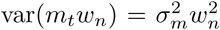 - see also Churchland et al. (2011)). Increasing the variance of the modulator leads to an increase in both neural variability and mean firing rate, due to the rectifying nonlinearity *g*(). As these two effects have opposing consequences for decoding, we chose to keep the mean firing rate constant as we varied modulator strength in our simulations. In the appendix we provide a more detailed examination of the effects of mean and variance on decoding (see Appendix A3). Fig. 6 shows the effect of inactive neurons on decoding performance. Here we only learn decoding weights for active neurons, from the two classes – task-informative (red) and task-uninformative neurons (black), in order to simplify and avoid unnecessary degrees of freedom. We thereby reduce the decoding problem to finding task-informative neurons among the active neurons. We set randomly varying baseline firing rates matched for informative and uninformative neurons.

**Table 1:** Knowledge assumed by each of the five decoders (Modulator Conditioned ML, Modulator Marginalized ML, Modulator Guided, Rate Guided, Sign Only - see text for details). Last column gives dimensionality of known variables, with *N* indicating the number of neurons in the population, and *T* the number of time points.

| Decoder | Stimulus response knowledge | Modulation knowledge | Degrees of freedom |
| --- | --- | --- | --- |
| MC-ML | $\lambda_n(s)$ (modulator-conditioned) | $m_t, w_n$ | $2N+N+T$ |
| MM-ML | $\lambda_n^*(s)$ (modulator-marginalized) | $\sigma_m, w_n$ | $2N+N+1$ |
| MG | none | $m_t$ | $T$ |
| RG | none | none | $0$ |
| SO | none | none | $0$ |

We computed decoding accuracy (proportion of correct discrimination choices) for a range of different modulator strengths, *σ*_*m*_. Results are shown in Fig. 7, plotted as accuracy (% correct) as a function of modulator strength (percentage of total spike count variance that can be attributed to the modulator). The MC-ML decoder gives a strict upper bound on decoding performance, as it assumes full knowledge of the encoding model. As the modulator variance *σ*_*m*_ increases, the performance of this decoder monotonically decreases, confirming that the added correlated noise is detrimental for encoding. For the encoding population tested here, the MM-ML decoder is nearly as good as the MC-ML decoder, although performance falls a bit faster with modulator variance. This is due to the use of a fixed threshold, that does not adjust to temporal fluctuations of the modulator. The rate-guided decoder performs only slightly above chance at all modulator levels, which confirms that the rate here does not contribute decoding-information that would allow differentiating between active-uninformative and informative neurons. Since inactive neurons are excluded from this simulation, the SO decoder performs at a similar level (compare to Fig. 6 and Appendix Fig. A1).

**Figure 7:**
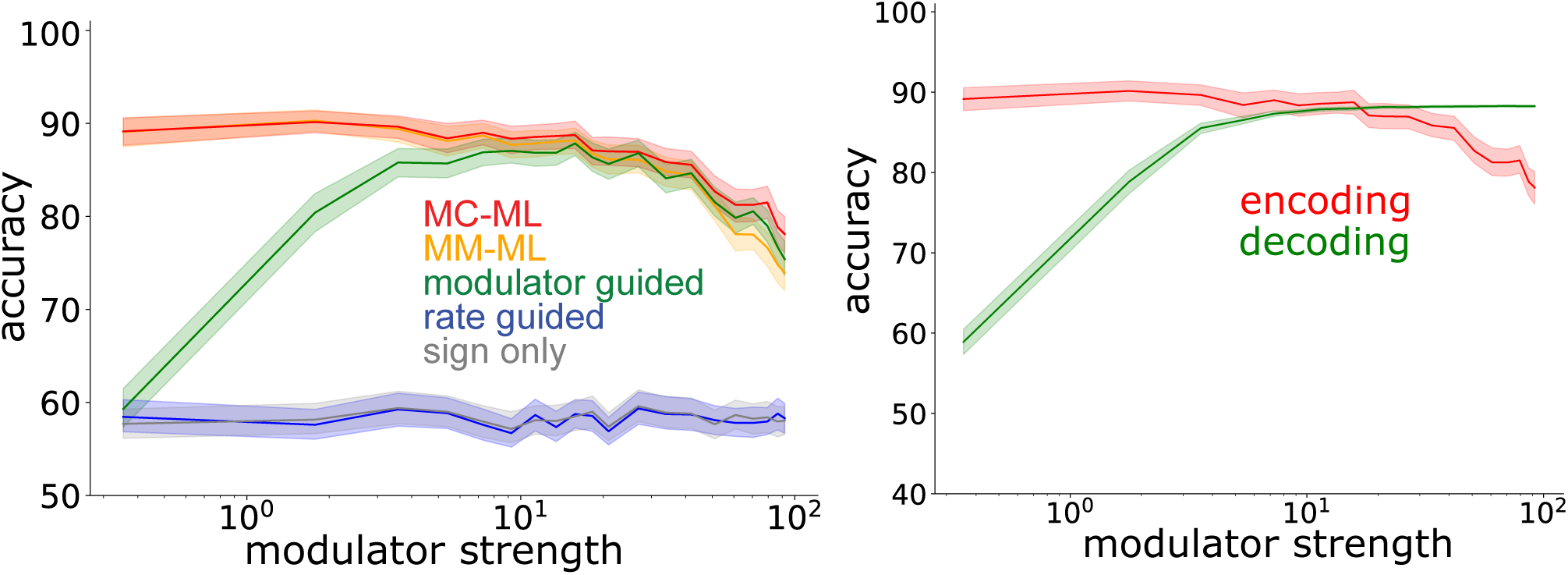
Left: Comparison of decoding accuracy of different decoders on simulated data. Curves indicate mean accuracy over all simulations, and its 95% confidence interval. Right: Increasing modulator strength has opposite effects on encoding and decoding accuracy.

The modulator-guided decoder exhibits a non-monotonic performance with increases in modulator strength. At low levels of modulation, performance increases with modulator strength - in this regime, the modulation allows the decoder to assign larger weights to the most informative cells. At higher levels, performance decreases with modulator strength, as with the ideal decoders, reflecting the corruption of the encoded signal. Hence, there exists an ideal level of modulation at which the MG decoder reaches its best performance, which is close to that of the ideal MC-ML decoder. Note also that since the MG decoder is capable of adjusting its threshold over time, depending on the modulator, it may outperform the MM-ML decoder (which uses a constant threshold) in the regime of high modulator strength.

In Figure 7 the two effects of the modulator are demonstrated separately by simulating activity in the same population and task-context at two stages; first to estimate decoding weights and second to read out the encoded activity. Noise from the modulator affects both stages differently. First, we can set decoding weights to be optimal and assess the effect of the modulator on encoding alone (second stage). As the modulator strength is increased encoding becomes noisier and the upper bound on possible information readout about the stimulus decreases (encoding in Fig. 7). Second, we use the MG decoder to detect informative neurons. We increase the modulator strength during the first stage of estimating decoding weights, leading to more accurate estimation of decoding weights, and keep the modulator strength constant during the second stage of readout. The % correct accuracy then reflects the effect of modulator strength on estimation of decoding weights without the impact on encoding. It is this trade-off between the impact on encoding and decoding that determines the maximum accuracy achieved by the modulator-guided decoder in Fig. 7 given varying degrees of modulator strength.

We then test the robustness of these results with respect to the percentage of informative encoding neurons, reflecting either task difficulty or intrinsic properties of the encoding population. We find that, as expected, increasing the percentage of informative neurons increases decoding accuracy for all decoders (see Appendix Fig. A3). As a result the RG decoder achieves reasonable decoding accuracy if more than half of the neurons are informative. The advantage of the MG decoder over the RG decoder is therefore strongest if the % informative neurons is small. This suggests that, within the experimentally-constrained parameter range, a modulator-guided decoding mechanism could indeed have a strong benefit over the simpler decoders.

## 4 Discussion

Flexible and robust decisions require dynamic routing of relevant sensory information in the brain, but the process by which this occurs is not understood. Here we have proposed a solution based on structured noise. We hypothesize that a functionally targeted stochastic modulator, as seen experimentally (Rabinowitz et al., 2015), could serve to dynamically label informative neurons, facilitating their readout, and thus enabling flexible and accurate decoding.

We showed that a modulator-guided linear decoder, in which weights are estimated through correlation of responses with the modulator, can achieve near-optimal performance. We investigated how parameters of the encoder (proportion of inactive neurons, and active but uninformative neurons) impact performance and found that these dictate a choice of modulator strength that best balances the disruptive effects of correlated noise on encoding against its positive effects for decoding. Importantly, performance is invariant to other parameter changes, such as size of the population and baseline firing rate, demonstrating the robustness of the modulation labeling scheme to circuit details.

While ideal observers set the functional form of the decoder, it remains unclear how to learn the weights in a biologically plausible way. Here we chose a simple heuristic that uses the inner product of spike count and modulator. There is a linear relationship between these heuristic estimates and the optimum, with no indication of systematic bias (Fig. A4). Moreover, the variance decreases with increasing modulator strength, suggesting that our heuristic provides a reasonable estimate of the optimal weights.

Historically, ideal observer models have ignored the presence of modulation, yet have provided good approximations of behavioral performance. Our MM-ML ideal observer provides an explanation for this incongruity: an experimenter that measures tuning functions by averaging neural responses in the presence of unaccounted-for modulation is effectively marginalizing over it. Optimal decoding weights derived from these estimates are in fact correct, but the use of a fixed decision threshold is suboptimal (it should vary in time with the modulator), which can results in temporally varying decision bias. For parameters in our simulation, this effect is relatively small (compare MC-ML and MM-ML in Fig. 7. Another source of bias, often ignored in previous ideal observer analyses, can arise from the selection of experimentally recorded neurons. This is generally biased towards active neurons, partly because low firing neurons are more likely to be overlooked, and partly because experimental stimuli are often optimized to drive the recorded population. While the signal may be concentrated in the recorded subpopulation, a downstream decoding area must also operate on the full (substantially larger) unrecorded population.

The need for flexibly routing information has been long recognized in the literature. One proposal relies on oscillations as a labeling mechanism (Singer, 1999). The idea is that the signal transmitted by sensory neurons is enhanced when their firing is synchronized. The “communication through coherence” theory (Akam and Kullmann, 2012) refines this idea in an encoding-decoding framework, in which a top-down oscillatory signal projects to both encoding neurons with the same feature selectivity, and to the decoding network that needs to read out from them. While oscillations in this theory play a similar role to our modulation scheme, there are several important distinctions. First, the oscillations in (Akam and Kullmann, 2012) are assumed to target feature-selective neurons, while we assume targeting of task-informative neurons. These could be the same for a detection task, but differ for a discrimination task (as in our simulations). Second, they assume a decoder with fixed threshold, which (as discussed above) is suboptimal. Third, they assume periodic oscillatory input, while we assume stochastic input that does not rely on any temporal structure (although note that our framework could also function with an oscillatory modulator). Finally, their framework describes a fixed labeling strategy, while ours is focused on flexible learning of task structure.

Our theory is agnostic to the nature of the modulator itself. Three important questions arise in this regard: 1) where is the modulatory signal generated, 2) by what mechanism does the modulator target encoding neurons in a task-specific manner, and 3) is the modulator signal available to the downstream decoder? Dynamic changes in noise correlations could arise through either local or top-down mechanisms. On the one hand (Bondy et al., 2018) suggests that a top-down source of the modulation changes pairwise correlations in V1 in a task-dependent manner. On the other hand, (Huang et al., 2019) show that detailed noise correlation structure can be produced locally. Changes in these locally produced correlations could be triggered via recruitment of inhibitory neurons (Middleton et al., 2012). In either case, it is not clear how task-specific neurons would be targeted. In the top-down case, at least for the orientation discrimination task considered here, the modulator could take advantage of the topological organization of sensory codes (for instance, orientation-specific columns in visual Area V1) to label clusters of task relevant neurons, without explicit knowledge of their individual tuning functions. The induced stochastic structure would then propagate bottom-up to higher areas, serving to label task-relevant neurons that are no longer spatially clustered. This would yield a hierarchy of modulator-based labeling across the sensory processing stages. Experimentally, we should find a signature of the same modulator across areas. Consistent with this, (Ruff and Cohen, 2016) show that the pairwise correlations between V1 and MT increase with behavioral performance, triggered by attending towards a stimulus.

If this kind of spatially localized modulation was indeed an organizing principle of neural activity, it would predict that only features that are represented in spatially localized at some stage in the brain can be flexibly decoded. If this was not the case for a particular task, then performance should be impacted, either in speed or accuracy. Hence, comparing performance in tasks that rely on features known to be spatially organized versus tasks with no spatial feature map could expose fundamentally different processing and learning strategies.

There are predictions that arise from our theory and do not depend on the source or detailed nature of the modulator. First, the strength of modulation is a key feature in successfully encoding task-relevant information (see Figure 7) and thus solving the class of discrimination tasks analyzed in our model. Consequently, the modulator strength (% total variance from modulator) should be non-monotonically related to the behavioral task-performance. Second, neurons that are most strongly modulated should be most influential for behavior in the task. This suggests that between two equally informative neurons the neuron that is more strongly modulated should have the higher choice probability.

While we have shown that modulator-guided decoding weights can be learned locally, it is still unclear how many training trials are required. While modulator-guided learning could be important to reach good performance after a small number of trials, it does not rule out that a different decoding strategy might be employed with additional experience. In this case, the strength of the modulator could be high in the beginning and gradually decrease throughout learning. Finally, future work needs to address the mechanistic implementation of learning via synaptic plasticity.

The lack of a biologically plausible theory of neural decoding strongly limits our understanding of neural computation, with direct impact on clinical applications such as the development of brain-machine interface (BMI) technology and the study of sensory and cognitive dysfunction. Moreover, flexible task-dependent decoding is unsolved in the field of machine learning, suggesting that there is a fundamental difference in how encoding-decoding is attempted by artificial neural networks and how it is solved by the brain. We’ve presented a theoretical solution to the identification problem of decoding based on targeted stochastic modulation, and a modulator-guided biologically plausible decoder. Our proposed theory serves as an important stepping stone towards understanding information routing in neural circuits, and lies computationally in between naive, inaccurate decoders and unrealistically all-knowing decoders.

## Appendix

### A1 Derivation of optimal decoders

Given the modulated Poisson model, *k*_*nt*_(*s, m*_*t*_) *∼* Poiss (*λ*_*n*_(*s*) exp(*w*_*n*_*m*_*t*_)), and assuming that the modulator *m*_*t*_ and the modulation weights *w*_*n*_ are known, the log probability of the stimulus *s* at time point *t* given spike counts *k*_*nt*_ of the whole population, *n* = {1, 2 … *N*}, becomes:

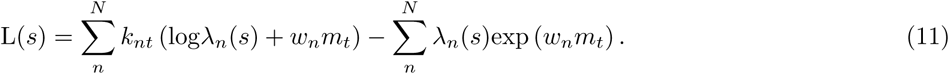

When discriminating between two stimuli *s* = {0, 1} under this model, the optimal decision is given by the sign of log-odds ratio, L(*s* = 1) − L(*s* = 0) ≷ 0, which translates into the following expression:

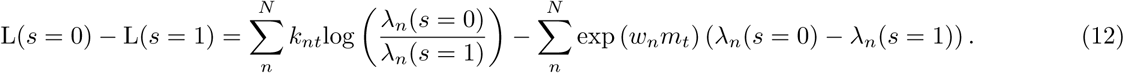

This corresponds to thresholding a weighted combination of individual neural responses, with optimal decoding weights:

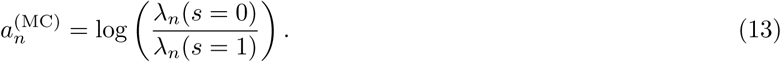

Since *m* is a known constant, it does not influence the decoding weights themselves, but merely changes the threshold. For the same reason, the optimal decoding weights remain unchanged whether the modulator is known (MC-ML), or whether its effects are marginalized over (MM-ML).

### A2 Inactive neurons

Figure 4 shows the 3 task-specific subpopulations into which encoding neurons can be divided. For computational efficiency, we excluded the subpopulation of inactive neurons from the simulations. Inclusion of those neurons affects mainly the SO decoder, for which the inactive neurons act as a source of noise. Although each one does not add much noise to the overall weighted sum, their contribution would be substantial in a real neural population, where they could constitute the majority of the cells. Figure A1 shows the changes in accuracy with increasing modulator strength for a population of 5000 neurons in which 4950 are largely inactive (grey population in Fig. 4). As expected, the performance of the SO decoder is strongly affected by the inactive neurons - its performance drops to chance level (50%).

### A3 Increasing spike count mean versus variance with increasing modulator strength

As we increase the modulator variance in simulation, it will not just introduce more noise but can also increase the mean response, depending on the chosen nonlinearity *g*(). In a biological network such an increase in mean could well be counteracted through adaptive gain-control (normalization), suggesting a constant mean. However, if this was not the case the expected increase in mean for our nonlinearity *g*(·) = exp(·) would be 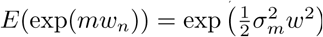. Such a boost in signal by itself would be beneficial for the encoding-decoding process, an example of stochastic resonance (McDonnell and Abbott, 2009). While in our main results we keep the mean constant, Fig. A2 shows the effect of increasing variance and mean separately or together through the modulator term. “modulator mean” hereby refers to the expected increase in firing rate as modulator variance is increased.

Figure A2 shows that increases in mean and variance have opposite effects on decoding. Increases in mean benefit encoding (see MC-ML) and heuristic decoders such as the rate-guided decoder (Fig. A2 left). In the simulation the modulator is a simple constant without variance (*m ∼ N* (*µ*_*m*_, 0)). This effect mixes with the detrimental effects of increasing noise variance (Fig. 7 in main text), leading to an overall mixed effect (Fig. A2 right).

### A4 Percentage of informative neurons

The decoding problem of identifying task-informative neurons is difficult in a task where only very few of the encoding neurons are actually informative. The percentage of informative neurons among the active population varies depending on the intrinsic tuning properties of the population (e.g., width of tuning curves), and extrinsic task properties (e.g., coarse vs fine discrimination task). The upper (MC-ML) and lower (rate-guided) bound on decoding accuracy will vary and consequently the relative advantage of a modulator-guided decoder depends on the nature of the task and the brain area. Varying the percentage of informative neurons in the population could provide a proxy for either changing intrinsic tuning properties or changing the discriminability of the two stimuli (i.e., “fine” vs. “coarse” discrimination task). Figure A3 shows decoder accuracy as a function of both modulator strength and percentage of informative neurons. We see that using the modulator for decoding offers an advantage especially if few neurons are informative.

### A5 Threshold for the modulator-guided decoder

The time-varying threshold for the MG-decoder (Eq. (10)) requires an estimate of the average change in activity as the stimulus changes Δ*λ*_*n*_(*s*). Given that we already have an estimate of which neurons are informative and which signed-group they belong to, this reduces to estimating the average stimulus-driven change in two subpopulations of informative neurons; Δ*λ*_+_(*s*) for the positively signed neurons and Δ*λ*_−_(*s*) for the negative ones.

### A6 Appendix figures

**Figure A1:**
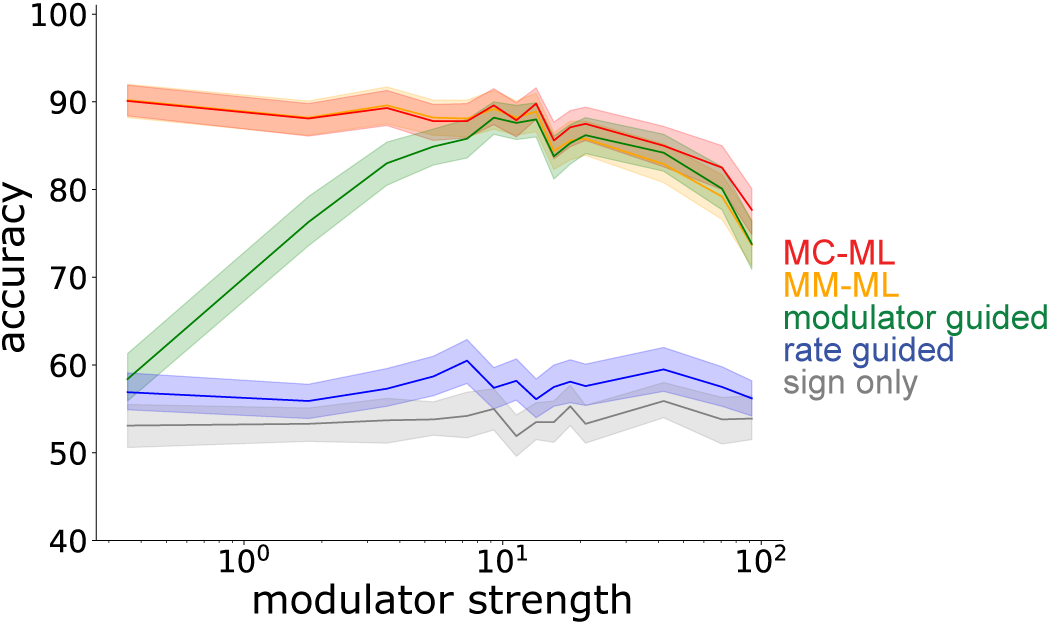
Comparison of decoding accuracy if inactive neurons are included. As in Figure 7 the decoding accuracy is assessed as the modulator strength is increased. However, instead of focusing exclusively on active neurons, we here include the population of inactive neurons. We see that only the SO decoder decreases performance while all other decoders are able to ignore the noise of inactive neurons.

**Figure A2:**
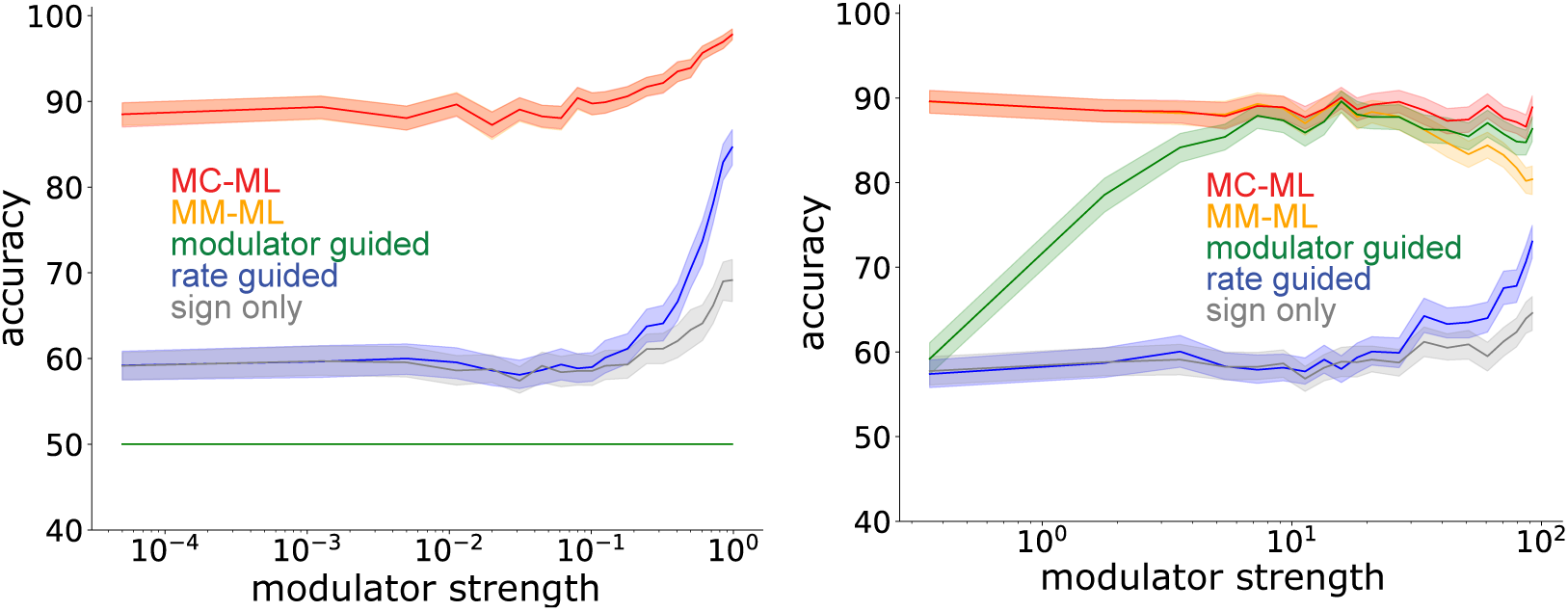
Mean spike count increase due to modulator. Increasing the variance of the modulator will increase both the variance of the spike count and the mean. While we kept the mean constant in the simulations for the main results, we assess the effect of mean and variance changes here. Left: Increase in mean only. The modulator is a constant input with increasing strength which increases the SNR and helps encoding, without introducing noise. There is no shared variability introduced by the modulator and a MG decoder cannot identify informative neurons. The MC-ML decoder and the MM-ML decoder perform equivalently. Right: Increase in both mean and variance. The modulator increases both variance and mean of the neurons that it modulates. The effect on decoding is a combination of these effects.

**Figure A3:**
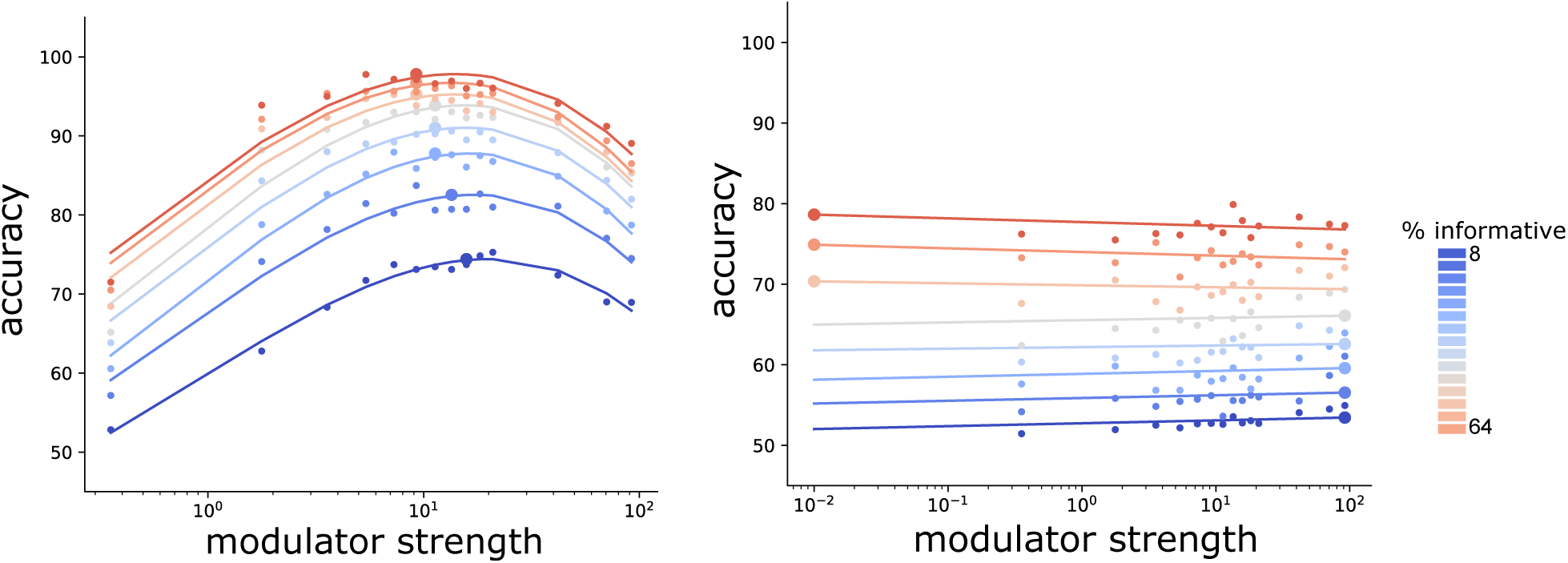
Percentage of informative neurons in encoding population. We increased the proportion informative neurons in the encoding population, thereby making the overall task easier. The performance of the MG decoder (left) has a maximum over modulator strength at any % informative. This maximum shifts towards smaller modulator strength values with larger % informative. The RG decoder is always outperformed by the MG decoder as long as there is still a substantial percentage of neurons that are active but uninformative.

**Figure A4:**
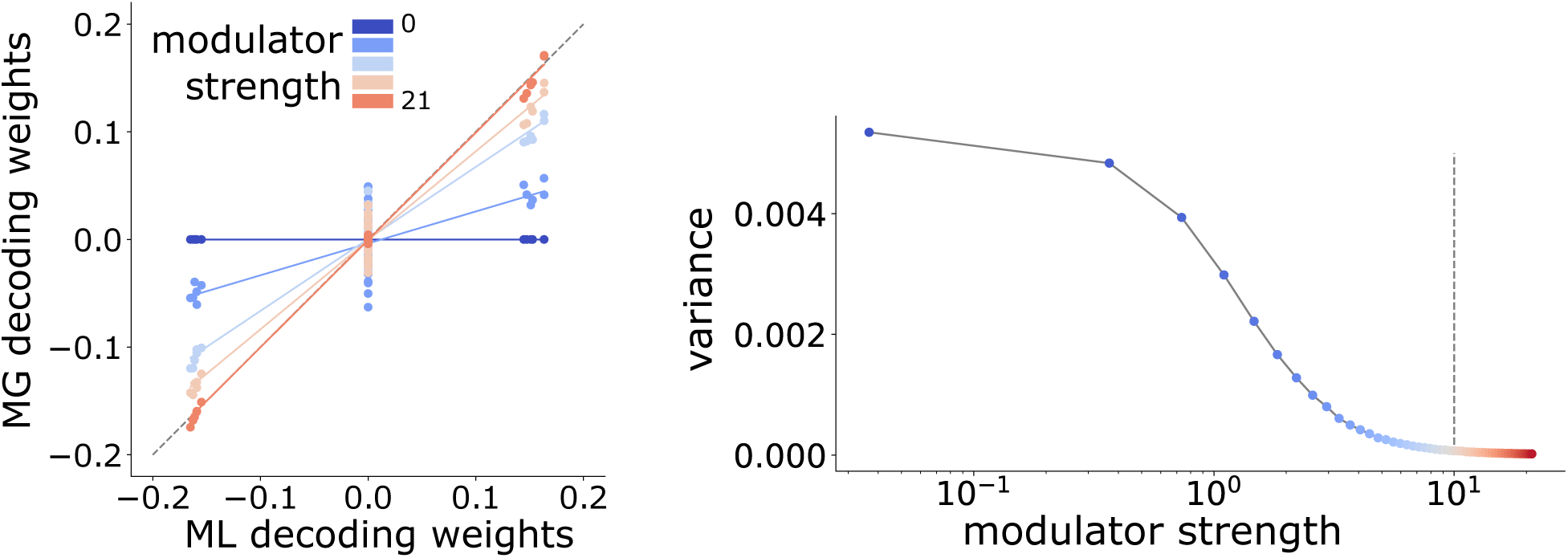
The same encoding population was simulated with a modulator of increasing strength. Decoding weights were computed optimally (ML decoding weights) and heuristically (MG decoding weights). Left) Mean of MG decoding weights at increasing strength of modulator. Relationship to true weights is linear and without an apparent bias. Right) The variance of the MG decoding weights decreases with increasing modulator strength. Grey line indicates the maximum performance of the MG decoder

## Footnotes

1 For brevity we use ‘decoder’ to refer to both the stimulus readout, and its corresponding optimal discriminator.

